# Identification of antibodies targeting the H3N2 hemagglutinin receptor binding site following vaccination of humans

**DOI:** 10.1101/675272

**Authors:** Seth J. Zost, Juhye Lee, Megan E. Gumina, Kaela Parkhouse, Carole Henry, Patrick C. Wilson, Jesse D. Bloom, Scott E. Hensley

## Abstract

Antibodies targeting the receptor binding site (RBS) of the influenza virus hemagglutinin (HA) protein are usually not broadly-reactive because their footprints are typically large and extend to nearby variable HA residues. Here, we identified several human H3N2 HA RBS-targeting monoclonal antibodies (mAbs) that were sensitive to substitutions in conventional antigenic sites and were not broadly-reactive. However, we also identified one H3N2 HA RBS-targeting mAb that was exceptionally broadly reactive despite being sensitive to substitutions in residues outside of the RBS. We determined that similar antibodies are present at measurable levels in the sera of some individuals but that they are inefficiently elicited by conventional vaccines. Our data indicate that some HA RBS-targeting antibodies can be surprisingly effective against variable viral strains even if they are somewhat sensitive to substitutions in HA residues adjacent to the RBS.

## INTRODUCTION

Influenza viruses continuously infect humans, in large part due to their ability to rapidly escape human immunity (Yewdell, 2011). Most neutralizing antibodies against influenza viruses target the globular head domains of hemagglutinin (HA) proteins and inhibit viral replication by blocking viral attachment. These types of antibodies often become ineffective after viruses acquire substitutions in epitopes within the HA globular head through a process called antigenic drift. As a result, seasonal influenza virus infections or vaccinations typically provide limited protection against antigenically drifted strains. New ‘universal’ vaccine antigens are currently being developed to elicit broadly-reactive antibodies against conserved epitopes in the HA receptor binding site (RBS) (Giles and Ross, 2011; Kanekiyo et al., 2019) as well as the HA stalk region (Impagliazzo et al., 2015; Krammer et al., 2013; Yassine et al., 2015).

New ‘universal’ vaccines that elicit antibodies against conserved epitopes in the HA RBS are attractive since antibodies against this region of HA directly block viral attachment and are highly neutralizing (Krause et al., 2011; Whittle et al., 2011). However, it is difficult to design appropriate vaccine antigens to elicit broadly neutralizing HA RBS-reactive antibodies because the surface area of most antibody footprints is larger than the narrow conserved RBS (Knossow and Skehel, 2006). The HA RBS is approximately 800 Å^2^ (Weis et al., 1988) whereas most antibody footprints are 1200-1500 Å^2^ (Amit et al., 1986).

Several broadly neutralizing antibodies that target conserved residues in the HA RBS have been identified (Ekiert et al., 2012; Krause et al., 2011; Lee et al., 2014; Lee et al., 2012; McCarthy et al., 2018; Schmidt et al., 2015b; Tsibane et al., 2012; Whittle et al., 2011; Winarski et al., 2015; Xu et al., 2013). These antibodies can arise from a number of V_H_ gene segments (McCarthy et al., 2018; Schmidt et al., 2015b). Most of these HA RBS-targeting antibodies bind through molecular mimicry, imitating the HA cellular receptor, sialic acid. Some of these broadly reactive antibodies make contact with conserved RBS residues through a shared dipeptide motif (Krause et al., 2011; Schmidt et al., 2015b; Whittle et al., 2011), while other antibodies insert a hydrophobic residue into the RBS (Xu et al., 2013). Most broadly reactive HA RBS-targeting antibodies possess atypically long HCDRs that allow the sialic acid-mimic motif of the antibody to guide into the conserved RBS while minimizing critical contacts to variable residues on the rim of the RBS (Ekiert et al., 2012; Lee et al., 2014; Whittle et al., 2011; Xu et al., 2013).

Here, we characterized the binding and neutralization characteristics of a large panel of anti-H3 human monoclonal antibodies (mAbs) that were isolated following seasonal influenza vaccination. Surprisingly, we found that a large proportion (>25%) of these mAbs targeted epitopes in the HA RBS. While most of these HA RBS-targeting mAbs were sensitive to substitutions in adjacent antigenic sites and were not broadly-reactive, we identified one mAb that maintained broad reactivity despite being moderately sensitive to substitutions at residues inside and outside of the RBS. We completed a series of experiments to further characterize this mAb and we determined that some individuals possess high levels of similar antibodies in polyclonal sera. These studies suggest that HA RBS antibodies are routinely elicited by vaccination and that some of these antibodies can be broadly reactive despite being sensitive to variation in residues adjacent to the conserved RBS.

## RESULTS

### Most vaccine-elicited human H3 mAbs target epitopes in HA globular head domain

We characterized 33 anti-H3 human mAbs that were isolated from 13 individuals vaccinated with the 2010-2011 trivalent seasonal influenza vaccine. First, we completed hemagglutination-inhibition (HAI) and micro-neutralization (MN) assays with the H3N2 component of the 2010-2011 vaccine, A/Victoria/210/2009, to determine if the mAbs prevent receptor binding and/or block virus infection *in vitro*. Twenty six out of 33 mAbs inhibited agglutination of the vaccine strain (Figure 1A), indicating that they likely targeted epitopes in the HA globular head domain. All HAI+ mAbs also neutralized the A/Victoria/210/2009 strain *in vitro* (Figure 1B). We identified 7 HAI-mAbs (Figure 1A), and we found that 2 of these mAbs neutralized virus *in vitro* while the remaining 5 were non-neutralizing (Figure 1B). Several HAI-mAbs inhibited binding of the HA stalk-reactive F49 mAb in competition assays (Supplemental Figure 1), suggesting that these mAbs targeted epitopes in lower regions of HA. These data are consistent with previous studies (Angeletti and Yewdell, 2018) that suggest that the majority of antibodies elicited by seasonal influenza vaccines target neutralizing epitopes on the HA globular head.

**Figure 1:**
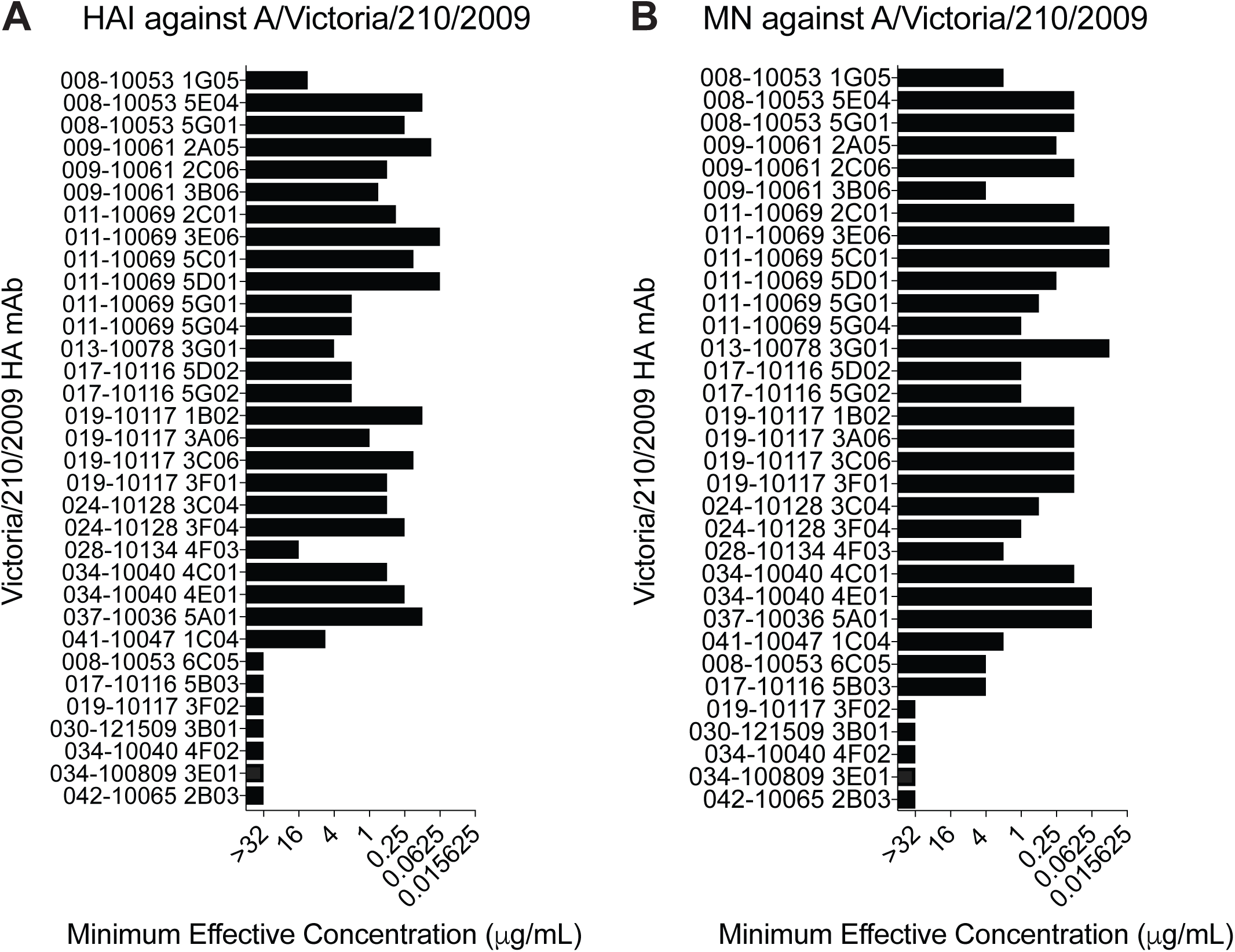
Hemagglutination-Inhibition and Micro-Neutralization Activities of mAbs. (A) Hemagglutination inhibition assays and (B) micro-neutralization assays were completed with each mAb and the A/Victoria/210/2009 vaccine strain. Titers shown are representative of two independent experiments.

### Most vaccine-elicited human H3 mAbs target HA antigenic site B

To map the footprints of each mAb, we measured binding to a panel of A/Victoria/210/2009 HAs that possessed different amino acid substitutions. For this, we created virus-like particles (VLPs) with A/Victoria/210/2009 HAs that possessed substitutions in classical antigenic sites (Koel et al., 2013; Wiley et al., 1981) and antigenic sites that have recently changed in naturally circulating human viral strains (Hadfield et al., 2018). Most of the substitutions in our panel were located in antigenic site A and B near the HA receptor binding site, but we also included several substitutions in epitopes in the lower part of HA head (Figure 2A). We also included HAs with substitutions in conserved residues within the RBS (Figure 2A), so that we could identify HA RBS-targeting mAbs. In total, we tested binding of all 33 mAbs using a panel of VLPs that expressed 25 different HAs in ELISAs (Figure 2B).

**Figure 2:**
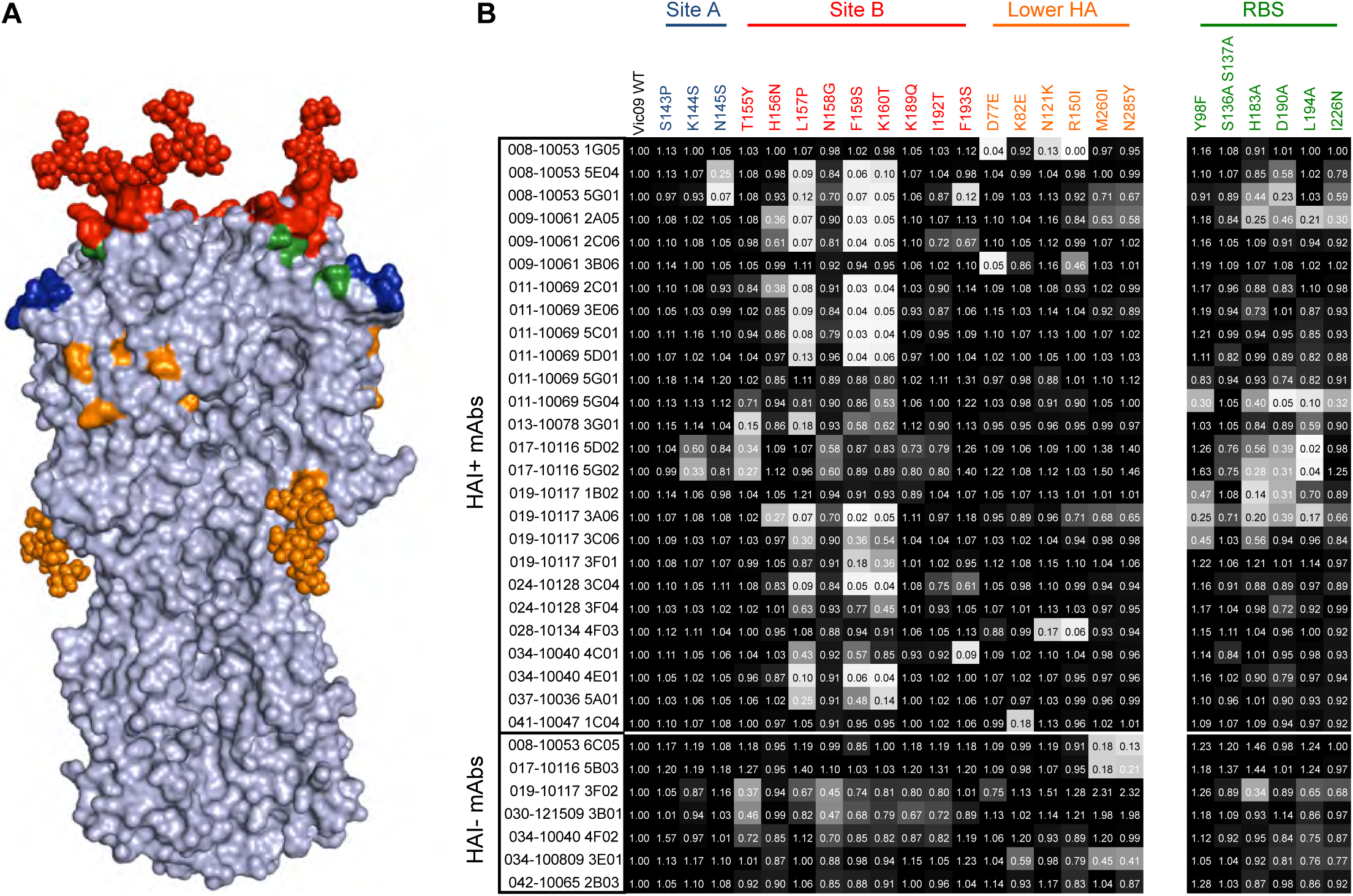
Antigenic Fine-Mapping of mAbs. Each mAb was tested for binding to a panel of HAs with different substitutions. (A) The location of each HA substitution tested is shown on the H3 structure. Most substitutions did not affect glycosylation, with the exception of the K160T substitution that results in the addition of a glycan at N158 (shown in red) and the N285Y substitution that results in the loss of a glycan (shown in orange). (B) ELISAs were completed using plates coated with VLPs bearing WT HA and HAs with different substitutions. Numbers in squares indicate fraction of binding relative to VLPs with A/Victoria/210/2009 WT HA. Colors of each square range from white (0% of binding to WT) to black (100% of binding to WT). Binding values are the average of two independent experiments.

The majority (73%) of HAI+ mAbs in our panel were sensitive to substitutions in HA antigenic site B. Mutations at residues 157, 159 and 160 of HA antigenic site B abrogated the binding of ∼61% of HAI+ mAbs, whereas substitutions at other antigenic site B residues (155, 156, 189, 192, 193) affected the binding of fewer mAbs. We identified several mAbs that were sensitive to substitutions in antigenic site A or epitopes lower on HA that were further from the receptor binding site. Only 3 mAbs in our panel were sensitive to substitutions in antigenic site A and 7 mAbs were sensitive to substitutions in residues lower on HA. These data suggest that antigenic site B is the major target of human neutralizing HA antibodies and demonstrate that single mutations near the RBS can abrogate the binding of most human mAbs.

While our data indicating that most of the mAbs in our panel are HA site B-specific are consistent with previous studies (Chambers et al., 2015; Popova et al., 2012; Zost et al., 2017), we were surprised to find that ∼1/3 of our HAI+ mAbs were sensitive to substitutions in conserved positions in the HA RBS (Figure 2B). Most of the mAbs in our panel that were sensitive to RBS mutations were also sensitive to site B mutations (Figure 2B). However, several mAbs that targeted the RBS were only moderately affected by site B mutations. For example, the 019-10117-3C06 mAb, which was moderately sensitive to RBS and site B substitutions, maintained partial binding to all of the mutant HAs that we tested. These data reveal that seasonal influenza vaccines unexpectedly elicit robust antibody responses targeting conserved residues within the HA RBS and that at least some of these antibodies can maintain partial binding to HAs that possess substitutions in conventional antigenic sites adjacent to the RBS.

### Identification of a HA RBS-targeting mAb with exceptional breadth

To assess the breadth of each mAb, we measured binding to HAs from H3N2 viruses isolated prior to and after the 2010-2011 season. As expected, most HAI-mAbs bound broadly to H3s isolated from 1968-2014 and two HAI-mAbs bound to both H3s and H1 (Figure. 3). In contrast, the majority of HAI+ mAbs bound to a narrow range of HAs from viruses that circulated from 2005-2012 (Figure 3). Most HAI+ mAbs failed to recognize an HA from a recent 2014 clade 3C.2a H3N2 strain (Chambers et al., 2015; Zost et al., 2017) that possesses an antigenically novel HA antigenic site B (Figure 3). Interestingly, 2 HAI+ mAbs (019-10117-3C06 and 028-10134-4F03) had exceptionally broad reactivity, binding to every H3 in our panel. One of these mAbs (028-10134-4F03) was sensitive to substitutions in residues 121 and 150 (Figure 2B) in the lower region of HA (Figure 2A). The second broadly reactive HAI+ mAb (019-10117-3C06) is particularly interesting because it is one of the HA RBS-targeting mAbs that we determined to be only moderately sensitive to substitutions in classic antigenic site B residues (Figure 2B). Despite having moderate reductions in binding to HAs with antigenic site B substitutions and RBS substitutions, the 019-10117-3C06 mAb maintained partial binding to every H3 in our panel, including the antigenically advanced 2014 clade 3C2.a H3N2 strain (Figure 3). The 019-10117-3C06 mAb originates from the IGHV1-69 germline and possesses a 19 amino acid HCDR3. The 019-10117-3C06 mAb also possesses a J_H_6 gene segment which has been previously reported as a common feature of mAbs recognizing the H1 RBS (Schmidt et al., 2015b).

**Figure 3:**
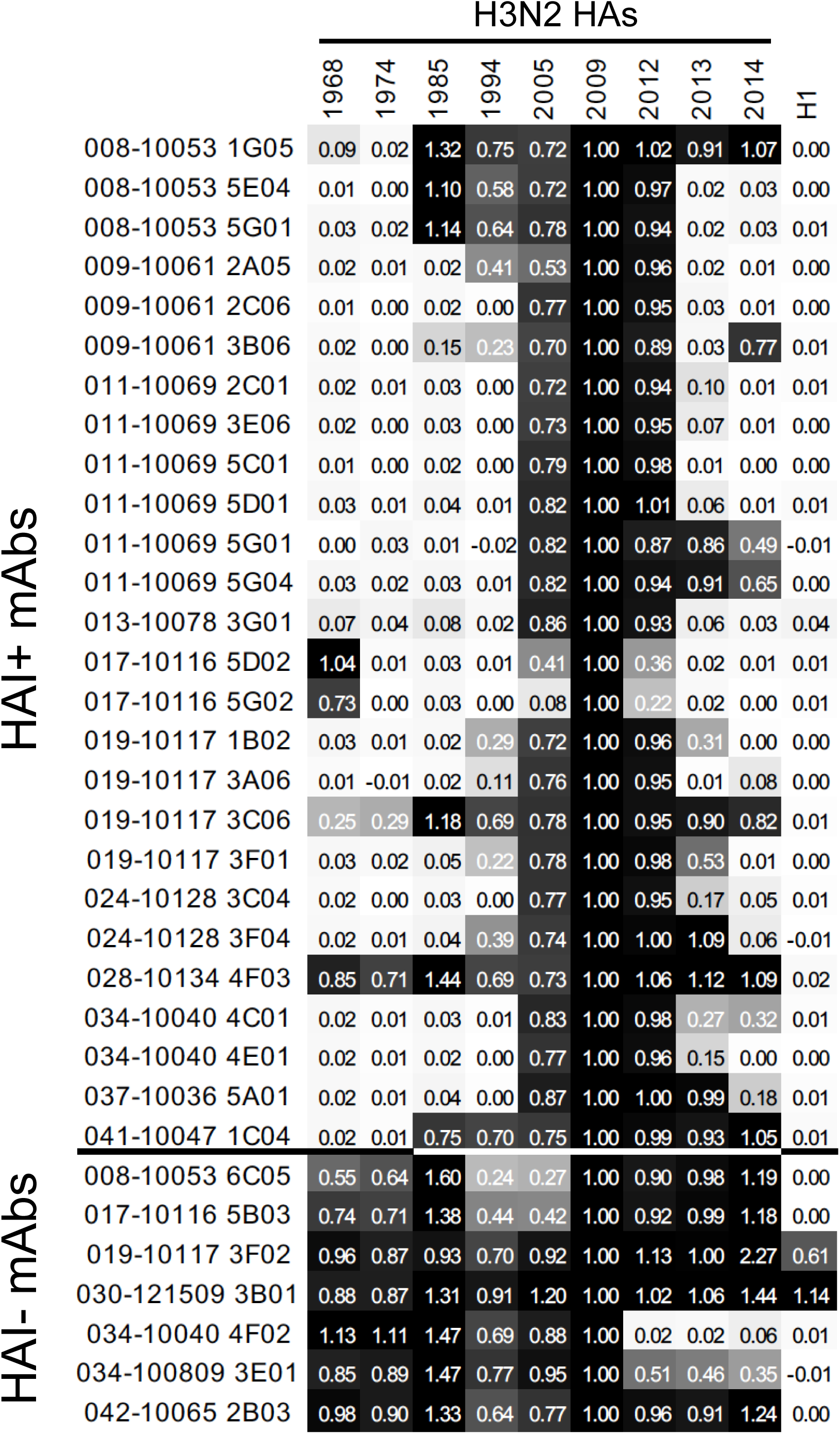
Binding of mAbs to Historical H3N2 Strains. ELISAs were completed with each mAb and a panel of historical H3N2 strains. Colors of each square range from white (0% of binding to WT) to black (100% of binding to WT). Binding values are the average of two independent experiments, while the value in each square indicates the fraction of binding relative to the A/Victoria/210/2009 vaccine strain.

### Characterization of a broadly reactive HA RBS-targeting mAb

We next completed a series of studies to further characterize the broadly reactive HA RBS-targeting 019-10117-3C06 mAb. First, we used a deep mutational scanning approach (Doud et al., 2017; Doud et al., 2018; Lee et al., 2018) to unbiasedly identify HA amino acid substitutions that could facilitate viral escape from this mAb. For these experiments we used a library of A/Perth/16/2009 HAs (which is antigenically similar to A/Victoria/210/2009) that possessed every possible single amino-acid substitution in HA and then we grew this virus library in the presence or absence of the 019-10117-3C06 mAb. For comparison, we completed parallel experiments where we grew the virus library in the presence of an HA antigenic site B mAb (024-10128-3C04) that did not have broad reactivity. As expected, the narrowly-reactive 024-10128-3C04 HA site B mAb selected viruses with substitutions in residues 159, 160, 192, and 193, which are located in HA antigenic site B (Figure 4A). Interestingly, the broadly reactive 019-10117-3C06 mAb also selected for viruses with HAs that possessed substitutions in antigenic site B (residues 159, 160, 193), as well as HAs that possessed substitutions in the adjacent antigenic site A (residue 145) (Figure 4B).

**Figure 4:**
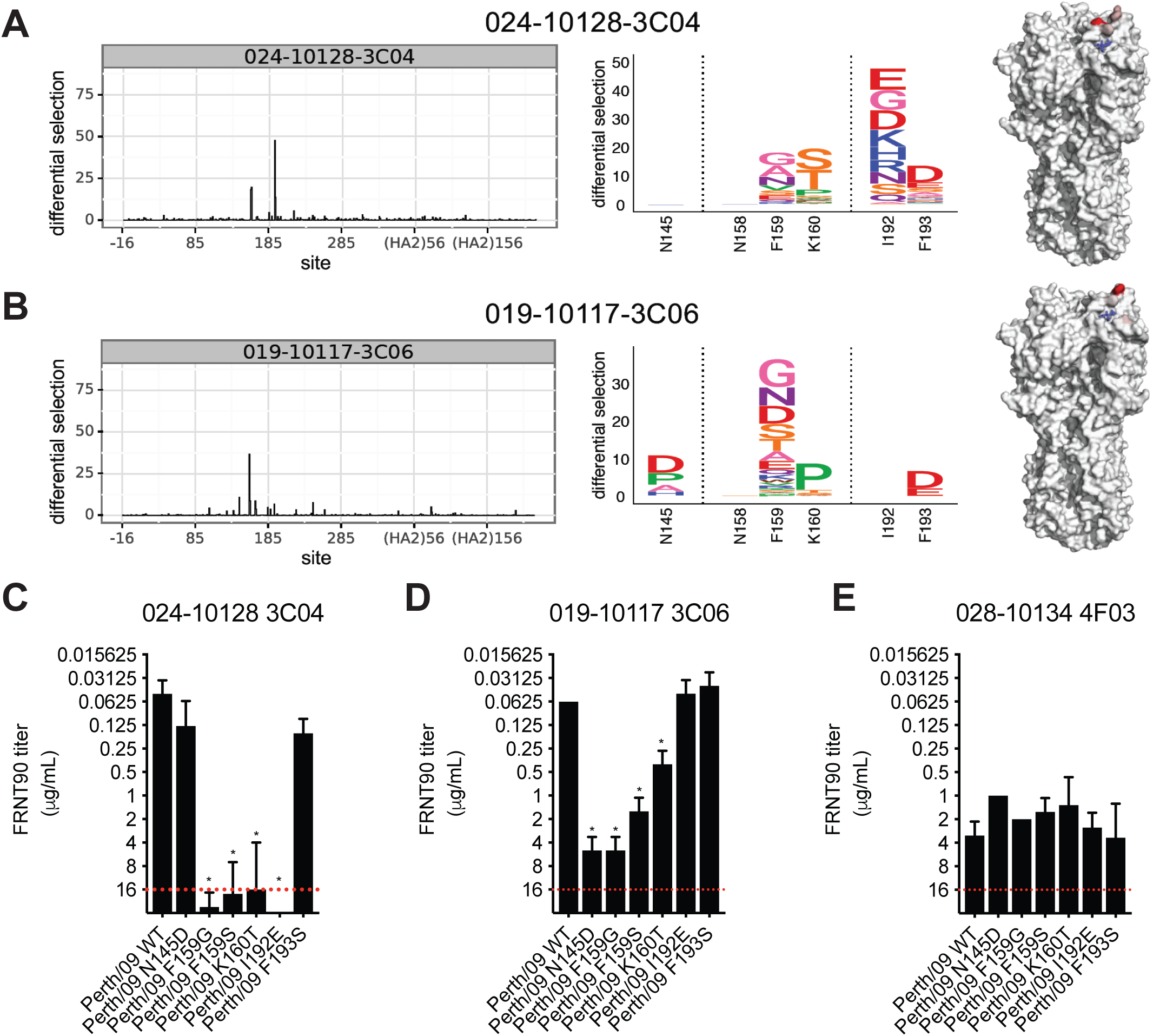
Mutational Antigenic Profiling of mAbs Targeting Antigenic Site B and the RBS. Deep mutational scanning experiments were completed to identify resistant viral fractions that survived after mAb selection. Logo plots showing the selection of amino acid substitutions and locations of these substitutions on the HA structure are show after selection with the 024-10128-3C04 mAb (A) and the 019-10117-3C06 mAb. Neutralization assays were completed with viruses that possessed several of the substitutions identified in deep mutational scanning experiments (C-E). A red dashed line indicates the limit of detection (each mAb was tested at a starting concentration of 16 ?g/mL). Neutralization titers shown are the geometric mean, and error bars denote the geometric standard deviation of three independent experiments. Statistical analyses of differences between FRNT titers against WT and mutant viruses were done using a one-way ANOVA with Dunnett’s test for multiple comparisons. *, p<0.05

In order to further characterize HA amino-acid substitutions identified in our deep mutational scanning experiments, we completed neutralization assays using viruses engineered to express A/Perth/16/2009 HAs with the N145D, F159G, F159S, K160T, or I192E substitutions. As expected, site B substitutions dramatically reduced neutralization of the narrowly-reactive 024-10128-3C04 mAb (Figure 4C). Substitutions at residues 145, 159, and 160 also reduced neutralization of the 019-10117-3C06 mAb, but importantly, all mutant viruses tested were still moderately neutralized by this mAb (Figure 4D). As a control, we also tested binding of the 028-10134-4F03 mAb, which binds lower on the HA head, against the 024-10128-3C04 and 019-10117-3C06 escape mutants. As expected, this mAb neutralized these mutants equivalently (Figure 4E). These data indicate that viruses can acquire HA substitutions that decrease neutralization of the broadly reactive 019-10117-3C06 mAb but that these substitutions do not completely escape from this antibody.

We hypothesized that the 019-10117-3C06 mAb is able to partially recognize viruses with HA antigenic site B substitutions by engaging conserved residues in the HA RBS. To test this hypothesis, we measured antibody binding to HAs that possessed a K160T HA substitution that introduces a glycosylation site in HA antigenic site B (Zost et al., 2017) with and without an additional Y98F substitution. HA residue 98 is located at the base of the RBS and interacts directly with sialic acid (Figure 2 and (Whittle et al., 2014)). Previous studies have shown that the Y98F substitution prevents HA binding to sialic acid without affecting the overall structure of HA (Bradley et al., 2011; Martín et al., 1998; Whittle et al., 2014). Consistent with our previous analyses (Figure 2), the 019-10117-3C06 mAb had moderate reductions in binding to HAs possessing either the K160T or the Y98F HA substitutions (Figure. 5A). Importantly, the 019-10117-3C06 mAb had dramatically reduced binding to HAs possessing both of these mutations (Figure 5A). As a control, we also tested binding of the narrow 024-10128-3C04 mAb to HAs possessing the K160T substitution with or without the Y98F substitution. Unlike the broadly reactive 019-10117-3C06 mAb, the narrow 024-10128-3C04 mAb failed to efficiently bind to HAs possessing K160T, with or without the Y98F substitution (Figure 5B). This suggests that partial binding of the 019-10117-3C06 mAb to HAs with antigenic site B substitutions is dependent on interactions with conserved residues in the RBS.

**Figure 5:**
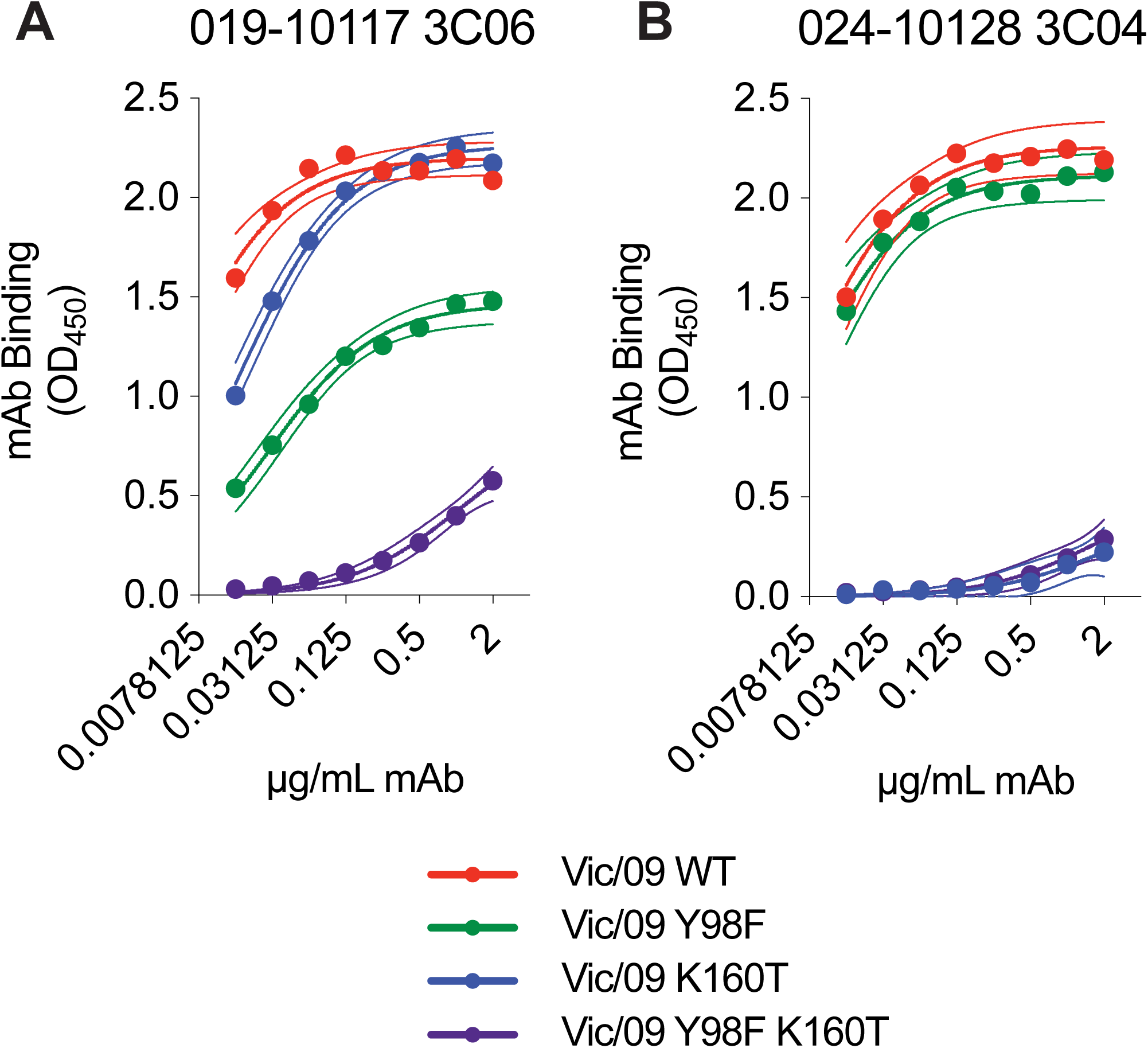
mAb 019-10117 3C06 Requires RBS Contacts for Cross-Reactivity. ELISAs were completed using the 019-10117 3C06 mAb (A) and the 024-10128-3C04 mAb (B) and plates coated with A/Victoria/210/2009 (Vic/09) HA VLPs with only Y98F, only K160T, or Y98F and K160T. ELISA binding curves from experimental triplicates are shown with one site – specific binding curves fit to the data (GraphPad Prism). Dashed lines represent the 95% confidence interval for each curve fit.

### Some individuals possess high levels of broadly-reactive HA RBS-targeting antibodies

We completed ELISAs to determine if HA RBS-targeting antibodies were present at high frequencies in the sera of donors pre- and post-vaccination. We tested sera from 28 individuals vaccinated during the 2010-2011 season, including 10 of the 13 donors that were used to generate mAbs. We tested sera antibody binding to ELISAs coated with A/Victoria/210/2009 HA, A/Victoria/210/2009 HA with a Y98F RBS substitution, A/Hong Kong/4801/2014 HA (a drifted strain with HA antigenic site B mutations), and A/Hong Kong/4801/2014 HA with a Y98F RBS substitution. Most sera samples did not have reduced antibody reactivity to HAs that were engineered to express the Y98F HA substitution, suggesting that the majority of antibodies in the serum of these vaccinated individuals were not directed against conserved residues of the RBS. However, serum antibodies from one donor (019-10117) had dramatically reduced binding to HAs with the Y98F substitution (Figure 6). Notably, the broadly reactive 019-10117-3C06 HA RBS-targeting mAb was derived from this same donor. This donor possessed Y98F-sensitive antibodies both prior to and after vaccination (Figure 6). Just like the 019-10117-3C06 mAb, polyclonal serum antibodies from this donor partially bound to the drifted A/Hong Kong/4801/2014 HA and binding of these antibodies was reduced by the Y98F HA substitution.

**Figure 6:**
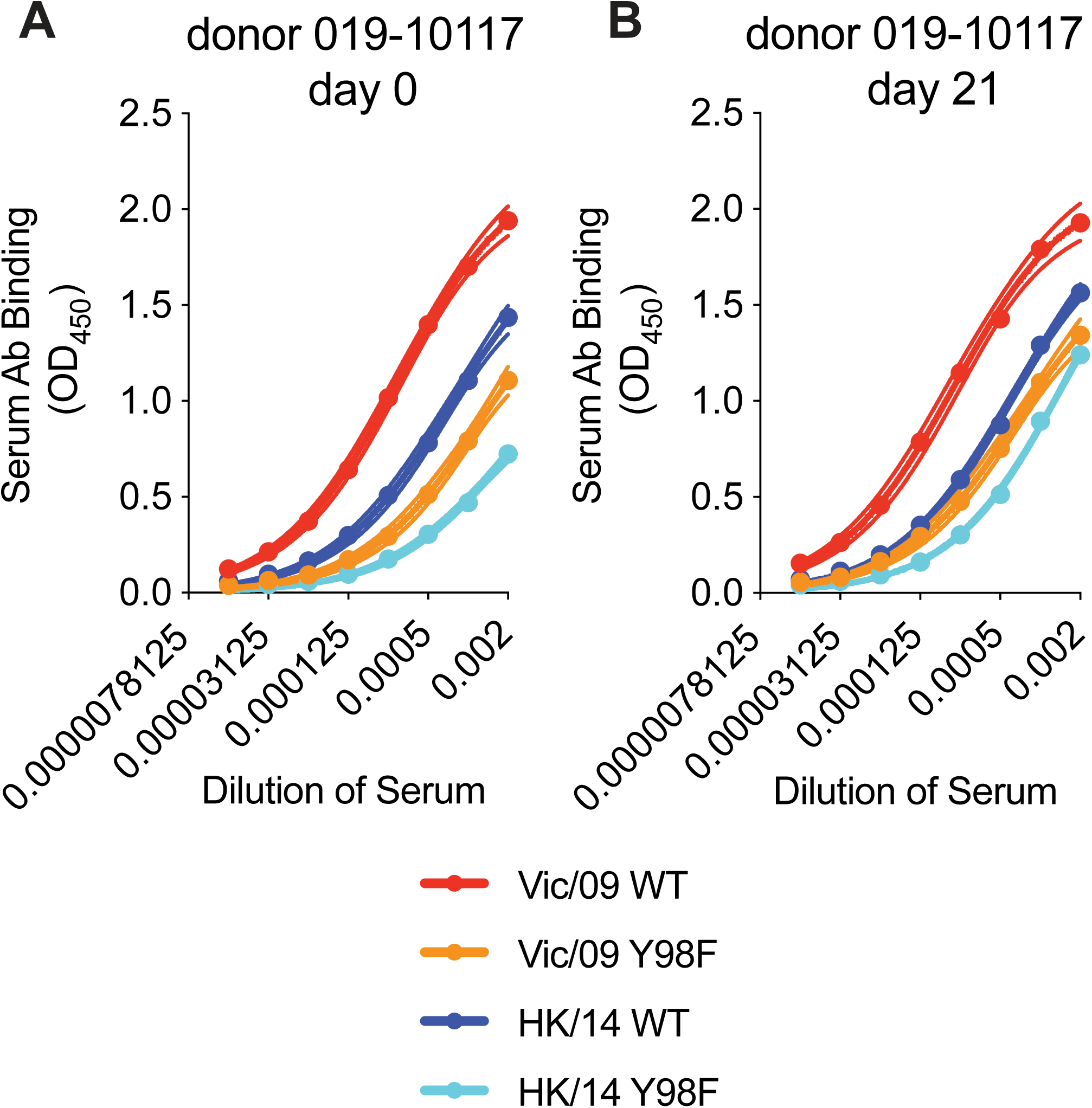
Rare individuals have high levels of RBS-targeting antibodies in serum. ELISAs were completed with serum from Donor 019-10117 with plates coated with A/Victoria/210/2009-WT HA (Vic/09 WT), A/Hong Kong/4801/2014-WT HA (HK/14 WT), and Y98F HA RBS mutants of both strains (Vic/09 Y98F, HK/14 Y98F). ELISAs were completed with serum collected prior to vaccination (day 0) (A) or day 21 following vaccination (B). Serum antibodies collected pre- and post-vaccination from 019-10117 exhibited reduced binding to A/Victoria/210/2009 HA with the Y98F substitution and the A/Hong Kong/4801/2014 HA with the Y98F substitution. Dashed lines represent the 95% confidence interval for each one site – specific binding curve fit.

### HA RBS-targeting antibodies are likely important in years with seasonal influenza vaccine mismatches

It is possible that HA RBS-directed antibodies are an important part of polyclonal neutralizing antibody responses during influenza seasons in which there are large antigenic mismatches between vaccine strains and circulating strains. It has been historically difficult to quantify levels of neutralizing HA RBS-directed antibodies in polyclonal sera. The main problem is that neutralization assays cannot be completed with HAs that have RBS substitutions since these substitutions often abrogate HA-sialic acid binding. To circumvent this problem, we developed an absorption-based approach to fractionate human serum samples. For these assays, we incubated serum antibodies with 293F cells expressing HAs with or without the Y98F substitution and then we completed *in vitro* neutralization assays with HA-absorbed serum fractions. In these assays, antibodies that are sensitive to the Y98F HA substitution are not absorbed by 293F cells that express the Y98F HA.

As a proof of principle, we first tested the broadly-reactive 019-10117-3C06 HA RBS-targeting mAb in this assay. We also included the broadly-reactive 041-10047-1C04 mAb that does not make contact with the HA RBS. For these experiments we tested binding and neutralization of the antigenically advanced A/Hong Kong/4801/2014 viral strain. Absorptions with 293F cells expressing the wild-type A/Hong Kong/4801/2014 HA completely removed both antibodies (Figure 7A). Conversely, absorptions with 293F cells expressing A/Hong Kong/4801/2014 HA with the Y98F substitution removed the control 041-10047-1C04 mAb but did not remove the HA RBS-targeting 019-10117-3C06 mAb (Figure 7A).

**Figure 7:**
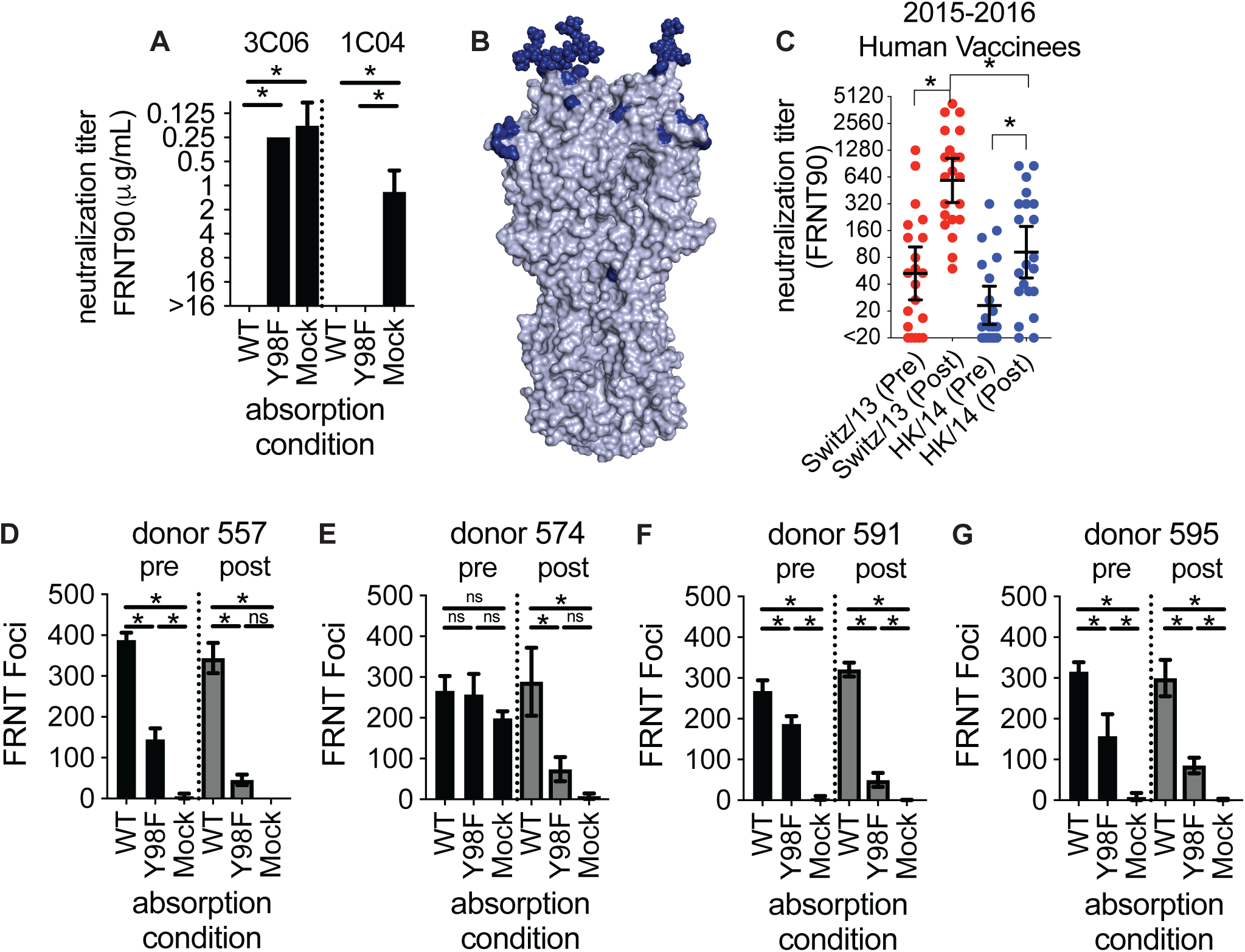
RBS Antibodies Contribute to Neutralizing Titers Against an Antigenically-Mismatched Strain. (A) Absorption assays were completed with the 019-10117-3C06 (abbreviated 3C06) and 041-10047-1C04 (abbreviated 1C04) mAbs. The A/Hong Kong/4801/2014-WT HA depleted both mAbs, while absorption with A/Hong Kong/4801/2014-Y98F HA did not deplete the 019-10117 3C06 mAb. Titers shown are the geometric mean of three independent experiments, and error bars show the geometric SD. (B) Residue differences between the HA of the 2015-2016 vaccine strain (A/Switzerland/9715293/2013) and circulating strain (A/Hong Kong/4801/2014) are shown in dark blue on the H3 structure. (C) Neutralization assays were completed with serum collected pre- and post-vaccination from individuals receiving the 2015-2016 seasonal influenza vaccine. Assays were completed with the vaccine strain (A/Switzerland/9715293/2013) and circulating strain (A/Hong Kong/4801/2014). Titers shown are the geometric mean of three independent experiments. Black lines indicate geometric mean and geometric 95% CI for each group, and statistical comparisons were made using the nonparametric Friedman test with correction for multiple comparisons. Statistically significant differences between groups are noted (*, p<0.05). (D-G) Absorption assays were completed with serum from individuals pre- and post-vaccination and neutralization of the A/Hong Kong/4801-WT virus was measured using an FRNT assay. Sera were absorbed with 293F cells expressing A/Hong Kong/4801-WT HA (WT), A/Hong Kong/4801 HA with a Y98F substitution (Y98F), or no HA (mock). Data are expressed as number of foci after absorbing a 1:80 dilution of each serum sample. Error bars show the mean number of foci +/- SD for three independent experiments. *, p<0.05

We next characterized serum antibodies in 21 individuals that received the 2015-2016 seasonal vaccine. We studied antibody responses elicited against the 2015-2016 vaccine because the H3N2 component of this vaccine was severely mismatched compared to A/Hong Kong/2014-like H3N2 viruses that circulated that season (Figure 7B). We first completed standard neutralization assays using the A/Switzerland/9715293/2013 vaccine strain and the antigenically distinct A/Hong Kong/4801/2014 virus. As expected, vaccine-elicited antibodies did not neutralize the mismatched A/Hong Kong/4801/2014 virus as efficiently as the A/Switzerland/9715293/2013 vaccine strain (Figure 7C). However, the A/Switzerland/9715293/2013 vaccine strain did boost serum antibody responses against A/Hong Kong/4801/2014 in some individuals. In order to determine if these cross-reactive antibodies were targeting conserved residues of the HA RBS, we completed absorption fractionation assays using serum from the same 21 vaccinated donors. For these experiments, we quantified the fraction of A/Hong Kong/4801/2014-reactive antibodies that were sensitive to the HA Y98F substitution. We detected Y98F HA-sensitive A/Hong Kong/4801/2014-reactive antibodies in 4 of 21 vaccinated donors (Figure 7D-G). We found that absorption with A/Hong Kong/4801/2014 WT HA depleted serum neutralizing antibodies, while absorption with A/Hong Kong/4801/2014HA-Y98F left an absorption-resistant fraction of neutralizing antibodies. Interestingly, some of these individuals had detectable RBS-targeting antibodies present prior to vaccination (Figure 7D,F,G). ELISA quantification confirmed that the antibodies left following A/Hong Kong/4801/2014HA-Y98F absorption bound to the A/Hong Kong/4801/2014 WT but not to the A/Hong Kong/4801/2014HA-Y98F HA, which indicates that this absorption-resistant fraction contained RBS-targeting antibodies (Supplemental Figure 2). These data suggest that cross-reactive HA RBS antibodies can be elicited by antigenically mismatched vaccines in some individuals, although this is not common with current egg-based vaccine formulations.

## DISCUSSION

A greater understanding of the specificity of anti-influenza virus antibody responses in humans is useful for rationally designing new universal influenza vaccine antigens. We began this study by antigenically charactering 33 H3 mAbs isolated from humans receiving a seasonal influenza vaccine. We found that the majority of these mAbs targeted epitopes in variable regions of the HA head. Some of these mAbs targeted conserved residues in the HA RBS but were not broadly reactive since they were also highly sensitive to HA substitutions in adjacent variable antigenic sites. However, we identified one HA RBS-targeting mAb that had exceptional breadth. This mAb (019-10117-3C06) was also moderately sensitive to HA substitutions in adjacent variable antigenic sites but was able to partially bind to antigenically drifted HAs.

Most HA RBS-targeting antibodies are not broadly reactive because their large binding footprints require contacts outside of the narrow RBS. However, several broadly reactive HA RBS-targeting antibodies have been identified (Ekiert et al., 2012; Krause et al., 2011; Lee et al., 2014; Lee et al., 2012; McCarthy et al., 2018; Schmidt et al., 2015b; Tsibane et al., 2012; Whittle et al., 2011; Winarski et al., 2015; Xu et al., 2013). A common feature of these broadly reactive HA RBS-targeting antibodies is that they all have relatively long HCDR3s, which allow them to minimize contacts on the rim of the RBS and maximize contacts with conserved RBS residues. Our study highlights that HA RBS-targeting antibodies can be broadly reactive even if they are moderately sensitive to substitutions in conventional antigenic sites near the RBS. The 019-10117-3C06 mAb from our study is clearly affected by substitutions in HA antigenic site B (Figure 2), but this antibody maintains binding to diverse H3 HAs (Figure 3) likely through multiple contacts with conserved residues in the RBS, which is facilitated by the antibody’s 19 amino acid HCDR3.

Our studies indicate that current vaccines do not efficiently elicit broadly reactive HA RBS-targeting antibodies in most individuals. We examined a cohort that received an antigenically mismatched vaccine, and although some of the donors mounted a cross-reactive antibody response, most of these cross-reactive antibodies were not binding to conserved residues in the HA RBS. Most conventional vaccine antigens are prepared in fertilized chicken eggs (Grohskopf et al., 2018) and contemporary egg-adapted H3N2 vaccine strains possess substitutions in or near the RBS which allow more efficient viral growth in chicken eggs (Wu et al., 2017; Zost et al., 2017). We speculate that vaccines that do not have adaptive mutations in the HA RBS might be better at eliciting antibodies targeting epitopes in the HA RBS of circulating viral strains. Future studies should determine if vaccine antigens that are not prepared in eggs are better able to elicit broadly reactive HA RBS-targeting antibodies.

The challenge, of course, is designing new vaccine antigens that are able to preferentially elicit antibodies like 019-10117-3C06. Recent work has sought to selectively elicit broadly-reactive HA head antibody responses through the use of “mosaic” nanoparticles that display antigenically diverse HA RBS domains on the same nanoparticle (Kanekiyo et al., 2019). This vaccination strategy might selectively activate naïve B cells targeting the HA RBS and might also selectively recall broadly-reactive memory B cells in secondary responses. One key challenge moving forward will to be determine if unique prior exposure histories facilitate the development of broadly reactive HA RBS-targeting antibodies. In our study, we identified some donors with very high levels of these antibodies in polyclonal sera. In the case of donor 019-10117, these antibodies were already at high levels in polyclonal sera prior to vaccination. Is there something genetically unique about donor 019-10117 or does that donor have a unique exposure history that gave rise to a B cell response highly focused on conserved residues within the HA RBS? While some studies have generated unmutated common ancestors and inferred the immunogenic stimuli for broadly-reactive antibody lineages targeting the HA RBS (McCarthy et al., 2018; Schmidt et al., 2015a), we know little about how prior immune history and repeated exposures influence the development of HA RBS antibodies. In the field of HIV, major efforts have been made to study antibody-virus co-evolution and the development of broadly neutralizing antibody specificities in chronically infected individuals, with the goal of identifying HIV envelope proteins that favor the development of broadly neutralizing antibody responses (Bonsignori et al., 2017; Landais et al., 2017; Rantalainen et al., 2018). Longitudinal studies in human cohorts could address similar questions for influenza viruses, with the potential to fill in gaps in our understanding of how antibody responses are elicited, recalled, and altered by infection and vaccination (Erbelding et al., 2018).

## ACKNOWLEDGMENTS

This work was supported by the National Institute of Allergy and Infectious Diseases (1R01AI113047, SEH; 1R01AI108686, SEH; CEIRS HHSN272201400005C, PCW, JDB, and SEH). Jesse D. Bloom and Scott E. Hensley hold an Investigators in the Pathogenesis of Infectious Disease Awards from the Burroughs Wellcome Fund.

## AUTHOR CONTRIBUTIONS

SJZ, JL, MEG, and KP completed experiments and analyzed data. CH and PCW provided plasmids that express heavy and light chains of each mAb in this study. JDB supervised mutational scanning experiments. SEH conceived the project, supervised the project, and analyzed data. SJZ and SEH wrote the manuscript with input from all authors.

## STAR METHODS

### Monoclonal Antibody Isolation and Purification

mAbs were isolated from human donors as previously described (Smith et al., 2009). Briefly, plasmablasts were single-cell sorted from peripheral blood mononuclear cells collected from donors seven days after vaccination with the 2010-2011 vaccine containing the H3N2 vaccine strain A/Victoria/210/2009. Single-cell RT-PCR was used to amplify V_H_ and V_L_ chains, which were cloned into human IgG expression vectors. mAbs were produced by transfecting 293T cells with plasmids encoding heavy and light chains and mAbs were purified using protein A/G magnetic beads.

### Hemagglutination-inhibition (HAI) Assays

mAbs were serially diluted twofold in a 96-well round-bottom plate in 50μL total volume of phosphate-buffered saline (PBS). After serial dilution, four agglutinating doses of virus in a total volume of 50 μL PBS were added to each well. Turkey erythrocytes (12.5 μL of a 2.5% [vol/vol] solution) were added and the sera, virus, and erythrocytes were gently mixed. After 1 hr at room temperature, plates were scanned and titers were determined as the lowest concentration of monoclonal antibody that fully inhibited agglutination. HAI assays were performed in duplicate on separate days.

### Microneutralization (MN) Assays

mAbs were serially diluted two-fold in round-bottom 96-well plates in 50μL serum-free Minimal Essential Medium (MEM). 50 μL of MEM containing 100 TCID_50_ of virus was added to serially diluted mAbs and the mAb-virus mixtures were incubated for 30 min at room temperature. Following incubation, the mAb-virus mixtures were added to confluent monolayers of Madin-Darby canine kidney (MDCK) cells in 96-well plates and incubated for 1 hr at 37°C. After incubation, the virus-antibody mixtures were removed and cells were washed with 180 μL MEM. After washing, serial dilutions of each mAb were added back to cell monolayers in infection media (MEM containing HEPES buffer, gentamycin, and 1 μg/mL TPCK-treated trypsin). The cells were incubated for 3 days and neutralization titers were determined as the lowest concentration of mAb that prevented cell death. MN assays were completed in duplicate on separate days.

### VLP Antigenic Mapping ELISAs

Point mutants of A/Victoria/210/2009 HA were generated in a codon-optimized HA gene by site directed mutagenesis. Virus-like particles (VLPs) were generated by transfecting 293T cells with each point mutant along with plasmids encoding HIV gag, the NA from A/Puerto Rico/8/1934, and a human-airway trypsin-like protease (HAT). Supernatants from transfected 293T cells were collected 3 days following transfection and were concentrated by centrifugation at 19,000 rpm (65,096 x g) in an SW-28 rotor using a 20% sucrose cushion. VLP pellets were resuspended in PBS and stored at 4°C. ELISA plates were coated with HA-normalized point-mutant VLPs diluted in PBS or just PBS as a background control and stored overnight at 4°C. The following day, plates were blocked with a 3% w/vol solution of bovine-serum albumin (BSA) in PBS for 2 hrs. After blocking, plates were washed five times with distilled water and two-fold serial dilutions of each mAb were added to plates in a 1% w/vol solution of BSA in PBS. After 2 hrs of incubation, plates were washed and a peroxidase-conjugated goat anti-human secondary antibody was added in a 1% w/vol solution of BSA in PBS. After incubation for 1 hr, plates were washed and 50μL of TMB substrate was added to each well. The TMB reaction was quenched by addition of 25 μL 250mM HCl and absorbance at 450nm was measured using a plate reader. In order to generate antigenic maps, one-site specific binding curves were fit to the data in GraphPad Prism software and the maximal binding (B_max_) was determined for each mAb. To generate antigenic maps from the ELISA data, we first selected the lowest mAb concentration that still gave at least 90% of the B_max_ signal. At this dilution, background signal was subtracted and signal for each point mutant was normalized to the A/Victoria/210/2009 WT HA VLP signal. Antigenic mapping ELISAs were conducted for each mAb in duplicate on separate days, and the resulting values were averaged and represented as a heatmap.

### Recombinant HA Production

Codon optimized HA genes for A/Hong Kong/4801/2014 WT and Y98F were cloned into expression vectors and the transmembrane was removed and replaced with the FoldOn trimerization domain from T4 fibritin, an AviTag site-specific biotinylation sequence, and a hexahistidine tag, as previously described (Whittle et al., 2014). Recombinant HAs were produced by transfecting 293F suspension cells with plasmids encoding HA. After four days, the supernatant was clarified by centrifugation and the HA proteins were purified by Ni-NTA affinity chromatography.

### Mutational Antigenic Profiling

We performed mutational antigenic profiling of mAbs 024-10128-3C04 and 019-10117-3C06 against A/Perth/16/2009 (H3N2) HA mutant virus libraries (Lee et al., 2018) using a previously described protocol (Doud et al., 2017). We selected two biological replicate libraries by incubating 1e6 TCID_50_ mutant viruses with neutralizing concentrations of antibody at 37°C for 1.5 hours. For antibody 024-10128-3C04, we neutralized mutant viruses with 0.1, 0.25, 0.2, 0.65, or 1 ug/ml antibody, and for 019-10117-3C06 we neutralized with 0.1, 0.2, 0.25, 0.55, 0.7, or 1 ug/ml antibody. We also included a mock selection condition where virus library was incubated with Influenza Growth Media (IGM, consisting of Opti-MEM supplemented with 0.01% heat-inactivated FBS, 0.3% BSA, 100 U of penicillin per milliliter, 100 ug of streptomycin per milliliter, and 100 ug of calcium chloride per milliliter). Following antibody incubation, we infected 2.5e5 MDCK-SIAT1-TMPRSS2 cells (Lee et al., 2018) with the virus-antibody mixture, then 2 hours post-infection aspirated off the inoculum, washed the cells with 1 mL PBS, then replaced the media with fresh IGM. Approximately 15 hours post-infection, we extracted, reverse-transcribed, and PCR amplified the viral RNA. We used the barcoded-subamplicon sequencing approach described in Lee et al., 2018 to deep sequence at high-accuracy. We then used dms_tools2 (v2.3.0) (Bloom, 2015) to analyze the deep sequencing results. The deep sequencing results are available on the NIH Sequence Read Archive under BioSample accessions SAMN10183083 (for the antibody-selected libraries) and SAMN10183146 (for the selection controls). The computer code for analyzing the data are at https://github.com/jbloomlab/Perth2009-HA_mAb_MAP

### Human Subjects and Serum Collection

All experiments involving humans were approved by the institutional review boards of the Wistar Institute, University of Pennsylvania, and the University of Chicago. Informed consent was obtained from all individuals. Experiments using deidentified human sera and mAbs were conducted at the University of Pennsylvania. For individuals from whom mAbs were isolated, serum was collected at the time of vaccination and 21 days post-vaccination. For individuals from the 2015-16 vaccination cohort, serum samples were collected at the time of vaccination and four weeks post-vaccination. For assays using foci-reduction neutralization tests, serum were treated with receptor-destroying enzyme (RDE) for 2 hrs at 37°C. Following treatment, the enzyme was heat-inactivated by incubation at 55°C.

### ELISAs with human sera

ELISA plates were coated the day prior with 0.5 μg/mL recombinant HAs (A/Hong Kong/4801/2014 WT, A/Hong Kong/4801/2014 Y98F, or a PBS background control) and blocked for 2 hrs on the day of the experiment with a 3% BSA in PBS solution. After washing the plates three times with wash buffer containing 0.5% Tween20 (vol/vol) in PBS (PBS-T), serially diluted serum samples were added to the ELISA plates and incubated for 2 hrs. After incubation, plates were washed three times with PBS-T and a peroxidase-conjugated goat anti-human secondary antibody diluted in 1% BSA in PBS was added. After 1 hr of incubation with the secondary antibody, plates were washed three times with PBS-T and 50 μL of a TMB substrate was added to each well. 25 μL of 250mM HCl was used to quench the reaction and the absorbance at 450nm was measured using a plate reader. Background signal at each dilution was subtracted for each serum sample and one site – specific binding curves were fit to the data using GraphPad Prism. Human sera ELISAs were performed in triplicate on separate days.

### Competition ELISAs

ELISA plates were coated the day prior with BPL-inactivated A/Hong Kong/1/1968 and blocked for 2 hrs with a 3% BSA in PBS solution. After washing the plates five times with distilled water, serial dilutions of the anti-H3 mouse mAb F49 or the control mouse mAb C179 in 1% BSA in PBS were added to the plate and incubated for 2 hrs at RT. After incubation, human mAbs were added directly to the plates at a fixed concentration in 1% BSA in PBS and incubated for 1 hr at RT. Plates were then washed five times with distilled water and peroxidase-conjugated anti-human secondary antibody was added and incubated for 1 hr. Following incubation with secondary antibodies, plates were washed five times with distilled water, TMB substrate was added, and the reaction was quenched with HCl. The absorbance at 450nm was quantified using a plate reader and competition at each dilution was normalized to the control mAb C179.

### Foci-Reduction Neutralization Tests (FRNTs)

RDE-treated serum samples were serially diluted in 96-well plates in a total volume of 50 μL. Approximately 200-300 focus-forming units of A/Hong Kong/4801/2014 WT virus in 50 μL were added to each well and the virus-absorbed sera mixture was incubated for 1 hr at room temperature. After incubation, the virus-sera mixture was added to confluent monolayers of MDCK-SIAT1 cells and incubated for 1 hr at 37°C. After incubation, cell monolayers were washed with 180μL serum-free MEM and an overlay medium containing HEPES, gentamycin, and 0.5% methylcellulose was added. The cell monolayers were incubated for 18 hrs, after which the overlay was removed and the cells were fixed at 4°C for 2 hrs using an aqueous solution of 4% paraformaldehyde (vol/vol). After fixation, cell monolayers were permeabilized using 0.5% Triton-X100 in PBS (vol/vol). After fixation and permeabilization, monolayers were blocked with a solution of 5% fat-free milk in PBS for 1 hr. After blocking, a mouse anti-nucleoprotein antibody was added in 5% milk/PBS for 1 hr. After the primary incubation, a peroxidase-conjugated goat anti-mouse secondary antibody in 5% milk/PBS was added for 1 hr. After incubation with the secondary antibody, monolayers were stained using a TMB substrate and foci were imaged and quantified using an ELISpot reader. For staining, plates were washed with distilled water between each step. Percentage of infection was determined relative to wells that did receive any serum or antibody. FRNT90 titer values are reported are the concentration of serum or mAb that reduced the numbers of foci by at least 90%.

### Absorption-Neutralization and Absorption-ELISA Assays

Two days prior to experiments, 293F suspension cells were transfected using 293fectin with plasmids expressing A/Hong Kong/4801/2014 WT HA, A/Hong Kong/4801/2014 Y98F HA, or a mock transfection control containing no plasmid or transfection reagent. On the day of the experiment, transfected cells were pelleted by centrifugation, washed twice with 293F media, and resuspended at the desired volume. In the case of absorption-neutralization assays, RDE-treated serum samples were diluted in 293F media at a dilution of 1:80 and split into three fractions for the three absorption conditions. An equivalent volume of 293F media containing approximately 8×10^6^ transfected cells/absorption reaction were added to each diluted serum sample and the samples were mixed by shaking for 1 hr at room temperature. After incubation, the cells were pelleted by centrifugation and the supernatant was transferred and re-centrifuged to clarify. Absorbed supernatant containing the sera was then serially diluted in 96-well round-bottom plates in serum-free MEM and FRNT assays were conducted as described. Absorption-neutralization experiments were completed in triplicate on separate days. In the case of absorption-ELISA assays, serum samples were diluted in 293F media at an initial dilution of 1:50 and split into three fractions for the three absorption conditions. Transfected cells were added and absorption of serum antibodies was carried out as described above. Following absorption, absorbed serum samples were serially diluted at a starting dilution of 1:500 (factoring in absorption volume) in 1% BSA w/vol in PBS and ELISAs were performed as described above.

For ELISA data, background antibody binding for each sample at each dilution was subtracted and one-site specific binding curves were fit to the data using GraphPad Prism software. The area under the curve (AUC) was calculated for each curve. For neutralization data, foci in positive control wells which did not receive any serum or antibody were used to adjust for variation between plates. Neutralization data are expressed as the number of foci remaining after absorbing a 1:80 dilution of serum. Assays were performed in triplicate on separate days. For both absorption-neutralization and absorption-ELISA experiments, the RBS mAb 019-10117 3C06 mAb and the 041-10047 1C04 mAb (which targets the lower HA head region) were initially diluted to a concentration of 32 μg/mL in 293F media prior to the addition of cells. For absorption-neutralization experiments, the starting concentration for each mAb absorption condition in the FRNT was 16 μg/mL. For absorption-ELISA experiments, the starting concentration for each mAb absorption condition in the ELISA was 3.2 μg/mL.

### Quantification and Statistical Analysis

Titer values for neutralization and ELISA experiments are reported as geometric mean and geometric SD, geometric mean and geometric 95% CI, or mean +/- SD as noted in each figure legend. For statistical analysis, the statistical tests used and the significance thresholds are described in the legend of each figure. All statistical analysis was performed in GraphPad Prism software.

