## Supplemental Figures for "Identification of antibodies targeting the H3N2 hemagglutinin receptor binding site following vaccination of humans"

**Zost et al. Supplemental Figure 1**

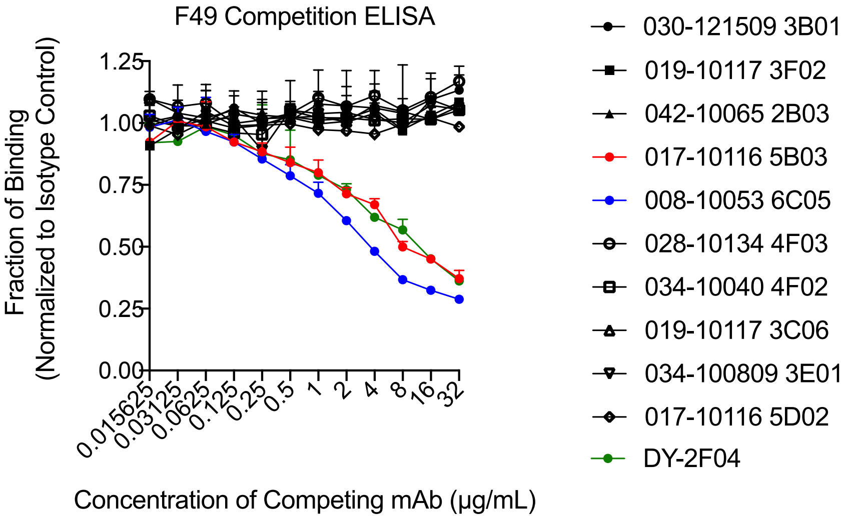


**Supplemental Figure 1: Competition ELISA with F49**
A mouse H3 stalk-specific mAb (F49) was used at varying concentrations to block binding of human mAbs to A/Hong Kong/1/1968 H3N2 virus. Competition at each dilution was normalized to the signal in the presence of the mock competitor C179, a group 1 HA mouse mAb that does not bind to H3 HA. Two human mAbs, 017-10116 5B03 (red) and 008-10053 6C05 (blue) compete with F49 for binding to H3 HA. DY-2F04, a previously described group 2 HA-stalk mAb, is included is a positive control (green).

### Zost et al. Supplemental Figure 2

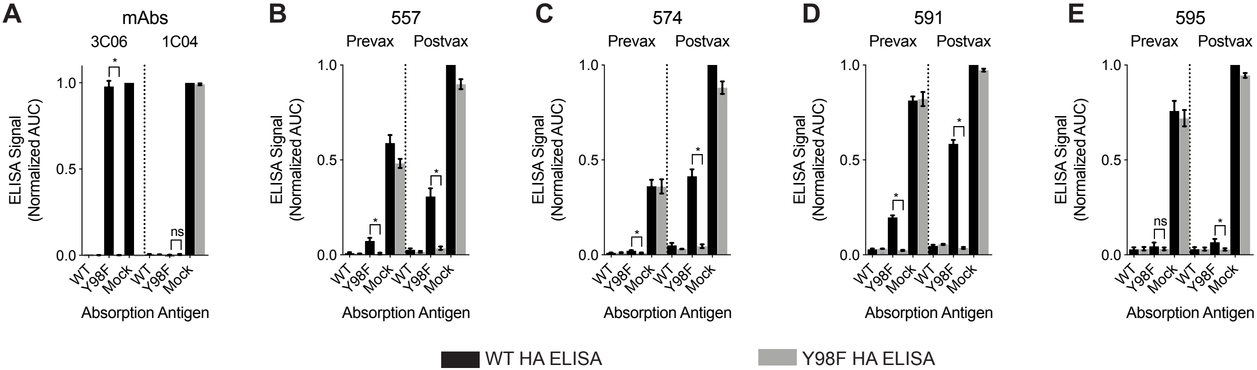


**Supplemental Figure 2: Serum antibody binding to HA before and after 2010-2011 vaccination**
Control mAb (A) and serum (B-E) ELISA titers against Hong Kong/4801/2014-WT HA and Hong Kong/4801/2014-Y98F HA (from Figure 7) after absorption with Hong Kong/4801/2014-WT HA, Hong Kong/4801/2014-Y98F, or cells only (mock). ELISA data are expressed as area under the curve (AUC) for each condition normalized to post-vaccination binding signal to Hong Kong/4801/2014-WT HA. Comparisons between WT and Y98F binding for the Y98F absorption condition were performed using an unpaired Student's t-test (*, $p<0.05$; ns = non-significant).
